# Lateral habenula mediates defensive responses only when threat and safety memories are in conflict

**DOI:** 10.1101/2020.11.12.378612

**Authors:** Geronimo Velazquez-Hernandez, Francisco Sotres-Bayon

## Abstract

Survival depends on the ability to adaptively react or execute actions based on previous aversive salient experiences. Although lateral habenula (LHb) activity has been broadly implicated in the regulation of aversively motivated responses, it is not clear under which conditions this brain structure is necessary to regulate defensive responses to a threat. To address this issue, we combined pharmacological inactivations with behavioral tasks that involve aversive and appetitive events and evaluated defensive responses in rats. We found that LHb pharmacological inactivation did not affect cued threat and extinction learning and memory, anxiety-like or reward-seeking behaviors. Surprisingly, we found that LHb inactivation abolished reactive defensive responses (tone-elicited freezing) only when threat (conditioning) and safety memories (extinction and latent inhibition) compete. Consistently, we found that LHb inactivation impaired active defensive responses (platform-mediated avoidance), thereby biasing choice behavior (between avoiding a threat or approaching food) towards reward-seeking responses. Together, our findings suggest that LHb activity mediates defensive responses only when guided by competing threat and safety memories, consequently revealing a previously uncharacterized role for LHb in experience-dependent emotional conflict.

## Introduction

In nature, survival in a dynamic environment requires animals to adaptively react or execute actions based on previous salient experiences (Elliot, 2008; Rangel et al., 2008; Namburi et al., 2016; Bravo-Rivera and Sotres-Bayon, 2020). In the laboratory, the regulation of reactive and active behaviors guided by stimuli that signal threat or safety and reward have been studied by using animal model combinations of aversive classical (Pavlovian) and appetitive instrumental conditioning (LeDoux, 2000; Cardinal et al., 2002). Auditory threat (fear) conditioning, in combination with lever pressing to obtain food, has been used to reliably evaluate defensive responses elicited after learning and memory of threat associations (tones paired with shocks) and its extinction (tones paired with omission of expected shocks) (Quirk et al., 2000; Sotres-Bayon et al., 2012). After extinction, the conditioned memory of threat and extinction memory of safety, coexist in the brain and compete for control of behavior (Quirk, 2007; Sotres-Bayon and Quirk, 2010; Lacagnina et al., 2019). Other experimental conditions that can be used to study the conflict between threat and safety memories is conditioned latent inhibition, which involves threat association of a tone that was previously associated with safety (Lingawi et al., 2017), as well as more recently developed tasks that involve the use of conflict choice behavior (Bravo-Rivera and Sotres-Bayon, 2020), such as platform-mediated avoidance (PMA) (Diehl et al., 2019). The use of these behavioral paradigms can help us characterize how the brain retrieves emotional memories to adaptively choose a defensive behavioral strategy.

The lateral habenula (LHb) has recently attracted much attention for its involvement in the regulation of a variety of behavioral responses triggered by aversive events (Hikosaka, 2010; Lammel et al., 2014; Proulx et al., 2014; Baker et al., 2016) including threat avoidance (Stamatakis and Stuber, 2012; Amo et al., 2014; Vincenz et al., 2017; Trusel et al., 2019; Stephenson-Jones et al., 2020), freezing (Martinez-Canabal et al., 2019), and choice behavior bias (Stopper and Floresco, 2014; Baker et al., 2017). The broad involvement of LHb in processing of aversive events is partly explained by its widely documented increase of activity in response to a variety of adverse stimuli, including those that were initially neutral but that have been conditioned to predict threatening events (Matsumoto and Hikosaka, 2009; Trusel et al., 2019). Yet, it is not clear if LHb is necessary to control threat-related behavioral responses ubiquitously. LHb lesions or pharmacological inactivations facilitated (Song et al., 2017), impaired (Thornton and Bradbury, 1989) or had no effect (Vale-Martinez et al., 1997; Shumake et al., 2010; Ilango et al., 2013) on threat avoidance. In addition, LHb contribution to another commonly evaluated defensive response, such as freezing, have been scarcely explored (Heldt and Ressler, 2006). Importantly, although LHb is predominantly involved with learning from negative events (Matsumoto and Hikosaka, 2009; Tian and Uchida, 2015), it has also been shown to signal positive value information (Stamatakis and Stuber, 2012). LHb neurons decrease their activity to positive stimuli, including those stimuli that have been conditioned to predict reward (Matsumoto and Hikosaka, 2009) and events that may signal safety through omission of threat (Li et al., 2019). Thus, given that LHb appears to simultaneously processes information about negative and positive events to guide motivated behaviors, it is possible that this brain region is critically involved under conditions that challenge individuals to convey information about opposing acquired values to adaptably control defensive responses.

To fully characterize LHb requirement in the control of defensive responses, we combined pharmacological inactivations with behavioral paradigms that involve threat stimuli, appetitive events and their competition. Surprisingly, we found that LHb activity is critical to promote reactive (freezing to a tone) and active (avoidance to a tone) defensive responses only when threat (conditioning) and safety (extinction and latent inhibition) memories are in conflict.

## Material and Methods

### Subjects

A total of 144 adult male Wistar rats (Institute of Cellular Physiology breeding colony) and weighing 260–280 g were housed in polyethylene cages and maintained on a standard 12 h light/dark cycle. Rats were housed individually, handled daily to diminish stress responses and received water *ad libitum* during all experiments. Rats were kept on a restricted diet (12 g/day) of standard laboratory rat chow to facilitate pressing a lever for sucrose pellets (Bioserv, Flemington, NJ). All manipulations and behavioral procedures were performed during the light phase between 7:00 and 19:00hrs. All procedures were approved by the Institutional Animal Care and Use Committee of the National Autonomous University of Mexico (UNAM) in compliance with the National Ministry of Health guidelines for the care of laboratory animals.

### Surgery

Rats were anesthetized with isoflurane inhalant gas and implanted with 23-gauge stainless steel guide cannulas (A-M Systems) targeting the lateral region of the habenula (−3.8 mm, AP, ±0.8 mm, −4.2 mm DV with respect to bregma). Cannulas were fixed with dental acrylic cement and anchored with two surgical screws placed on the skull. Stainless steel obturators were inserted into the guide cannulas to prevent clogging until infusions were made. The tips of the cannulas were aimed 0.8 mm above the target structure. After surgery, animals received food and water *ad libitum* for 7 d to allow full recovery before experiments.

### Behavior

Rats were trained to press a lever for sucrose pellets (Bioserv, Flemington, NJ) on a variable interval (VI) reinforcement schedule (VI-60 for conditioning tasks as in (Sierra-Mercado et al., 2011), and VI-30 for the avoidance task as in (Bravo-Rivera et al., 2014)). All rats received six lever-press training sessions until they reached a minimum rate of 12 presses per minute in their final session. Pressing to obtain food maintains a constant level of activity against which freezing, and avoidance can be reliably measured.

### Threat conditioning and extinction

Threat conditioning and extinction tasks were performed as previously described (Martinez-Canabal et al., 2019). Briefly, auditory threat conditioning and extinction was performed in standard operant chambers (Coulbourn Instruments, Allentown, PA) located inside sound-attenuating boxes (Med Associates, Burlington, Vermont, VT) in an isolated testing room. The floor of the operant chambers consisted of stainless-steel bars that could deliver electric foot-shocks. Between experiments shock grids, floor trays and walls were cleaned with soap and water. On day 1, rats were subjected to threat conditioning consisting of five tone presentations (30 s, 4 kHz, 75 dB) that co-terminated each one with a foot-shock (0.5 s, 0.6 mA). On day 2, after conditioning, rats received extinction training consisting of fifteen presentations of unreinforced tones. On day 3, rats were tested for memory retrieval with two tone-alone presentations. In all sessions, the interval between tones was variable with an averaged 2 minutes.

### Conditioned latent inhibition

On day 1, rats received latent inhibition training consisting of fifteen tone-alone presentations (30 s, 4 kHz, 75 dB). On day 2, rats were subjected to threat conditioning, consisting of five tones (30 s, 4 Hz, 75 dB) that co-terminated with a foot-shock (0.5 s, 0.6 mA). On day 3, rats were tested for memory retrieval with two tone-alone presentations. In all sessions, the interval between tones was variable with an average of 2 minutes.

### Platform-mediated avoidance task

PMA task was performed as previously described (Bravo-Rivera et al., 2014) using standard operant chambers (Coulbourn Instruments, Allentown, PA) located inside sound-attenuating boxes (Med Associates, Burlington, Vermont, VT) in an isolated testing room. The floor of the operant chambers consisted of stainless-steel bars that could deliver electric foot-shocks. Between experiments shock grids, floor trays and walls were cleaned with soap and water. Rats were conditioned to a tone (30 s, 4 kHz, 75 dB) co-terminating with a shock delivered through a grid floor (2 s, 0.4 mA). Rats received 9 tone–shock pairings each day for 10 days, with a variable interval between tones averaging 3 min. A safety square platform (14.0 cm per side) was located in the opposite corner of the lever to allowed rats to avoid the shocks. The availability of food on the side opposite of the platform motivated rats to leave the platform during the inter-trial interval, facilitating trial-by-trial assessment of avoidance. On day 11, rats received two tones to test for memory retrieval.

### Open field task

Rat locomotor activity in the open field (90 × 90 cm) was assessed by calculating distance travelled in the periphery and center (30 × 30 cm within) of the arena during 10 min. The overall distance traveled and number of entries to the center of the arena were used to assess locomotion and anxiety-like behavior, respectively.

### Lever pressing test

Rats were trained to press a lever to receive food pellets on a VI-60 schedule (as described above). Spontaneous presses per minute were calculated to assess reward-seeking behavior during 5 min.

### Drug infusion

We used GABA-A and GABA-B receptors agonists, muscimol and baclofen (M& B; Sigma-Aldrich) respectively, to enhance neuronal inhibition, thereby temporarily and reversibly inactivating the target structure. The inactivating drug cocktail was prepared on the day of the infusion using a filtered, physiological saline solution (SAL) as vehicle and infused 10 min before behavioral testing. M&B (50 ng of each drug / 0.2 μl per side) or SAL was infused at a rate of 0.2 μl delivered over 45s into the target structure. Cannulas were connected via polyethylene tubing to 10 μl syringes (Hamilton) driven by a programmable microinfusion pump (KD Scientific). Dosages and volumes of GABAergic agonists were based on a previous inactivation study targeting the same brain structure (Stopper and Floresco, 2014). After infusions, injectors were left in place for 2 min to allow the drug to diffuse.

### Histology

After all behavioral experiments, rats were deeply anesthetized with pentobarbital sodium (150 mg/kg, i.p.) to transcardially perfused with 0.9 % saline solution. Brains were then harvested and stored in a 10% formalin solution (Sigma-Aldrich) for at least 2 days. Then we switched the brains to a 30% sucrose solution for cryoprotection. Brains were cut in 45 μm-thick coronal sections using a cryostat (Leica, CM1520) and stained for Nissl bodies with cresyl violet and examined under a bright field microscope (Nikon, H550S) to verify cannula tip placements. Only animals with localized injector tips within LHb were included in statistical analysis; except for one experiment (LHb inactivation after extinction, Fig 2, in which analysis of data included animals where injector tips were localized within or outside the LHb (Hit M&B and Miss M&B groups, respectively).

### Data collection and analysis

All behavioral responses were recorded with digital video cameras (Logitech) and automatically analyzed with a commercial software (ANY-maze; Stoelting Co., IL, USA). Two measures of defensive responses were assessed throughout all experiments: 1) percent of time spent freezing and 2) suppression ratio. Freezing was defined as the absent of all movement, except respiration. The amount of time freezing to the tone was expressed as the percentage of the tone duration. We also measured time spent freezing before tone presentations (5 min), during context exposure (pretone freezing). There were little contextual-elicited defensive responses across all experiments (SAL and M&B groups), as evidenced by low levels of pretone freezing (all group averages < 12.03%). Matching of animal groups (experimental and controls) for similar levels of freezing responses before local drug infusions were done in all experiments (p= 0.95). A suppression ratio comparing pretone press rates with tone press rates was calculated as follows: (pretone - tone) / (pretone + tone). A value of 0 represents no suppression (low defensive response levels), whereas a value of +1 represents complete suppression of lever pressing (high defensive response levels). Avoidance responses were assessed by calculating percent rats that avoided the footshock by moving into to the platform during the last two seconds of tone presentations and time spent on platform before and after tone presentation. Groups were compared by using, when appropriate, unpaired Student’s two-tailed t tests, one-way or repeated-measures analysis of variance (ANOVA) followed by *post hoc* Fisher’s least significant difference test ((STATISTICA; StatSoft, Tulsa, OK) and Prism (GraphPad, La Jolla, CA)). Alpha was set at *p* = 0.05.

## Results

Although LHb activity has been broadly implicated in the regulation of aversively motivated behaviors (Thornton and Bradbury, 1989; Matsumoto and Hikosaka, 2009; Stamatakis and Stuber, 2012; Amo et al., 2014; Lecca et al., 2017; Song et al., 2017; Vincenz et al., 2017; Martinez-Canabal et al., 2019; Trusel et al., 2019), it is not always necessary for regulating defensive responses to a threat (Vale-Martinez et al., 1997; Heldt and Ressler, 2006; Shumake et al., 2010; Ilango et al., 2013; Song et al., 2017). To systematically evaluate the necessity of LHb under different aversive conditions, we combined pharmacological inactivations with behavioral tasks that involve aversive and appetitive motivations and evaluated defensive behavioral responses in rats.

### LHb inactivation does not affect threat or extinction learning and memory

To evaluate the requirement of LHb on threat learning and memory, we pharmacologically inactivated the LHb before auditory threat conditioning on day 1 (**Fig. 1**, ***A***_***1***_). We found that inactivation of LHb before threat conditioning did not affect threat acquisition or its retrieval (**Fig. 1**, ***A***_***2***_), as indicated by similar levels of defensive responses across groups (day 1, freezing: *F*_(1,10)_ = 0.71, *p* = 0.41; lever-press suppression ratio: SAL: *t*_(10)_ = −0.25, *p* = 0.80) and the first two-tone trial block of extinction training (day 2, freezing: *t*_(10)_ = 0.03, *p* = 0.96; lever-press suppression ratio: *t*_(10)_ = 0.26, *p* = 0.79), respectively. Also, LHb inactivation prior to threat conditioning did not affect extinction acquisition one day later or its retrieval two days later, as indicated by similar levels of defensive responses across groups in extinction training (day 2, freezing: *F*_(6,60)_ = 0.61, *p* = 0.71; lever-press suppression ratio: *F*_(6,60)_ = 0.52, *p* = 0.78) and in the memory test (day 3, freezing: *t*_(10)_ = −0.22, *p* = 0.82; lever-press suppression ratio: *t*_(10)_ = 1.12, *p* = 0.28). Thus, consistent with previous reports (Heldt and Ressler, 2006; Song et al., 2017), our findings suggest that LHb is not necessary for threat learning and memory.

**Figure 1.**
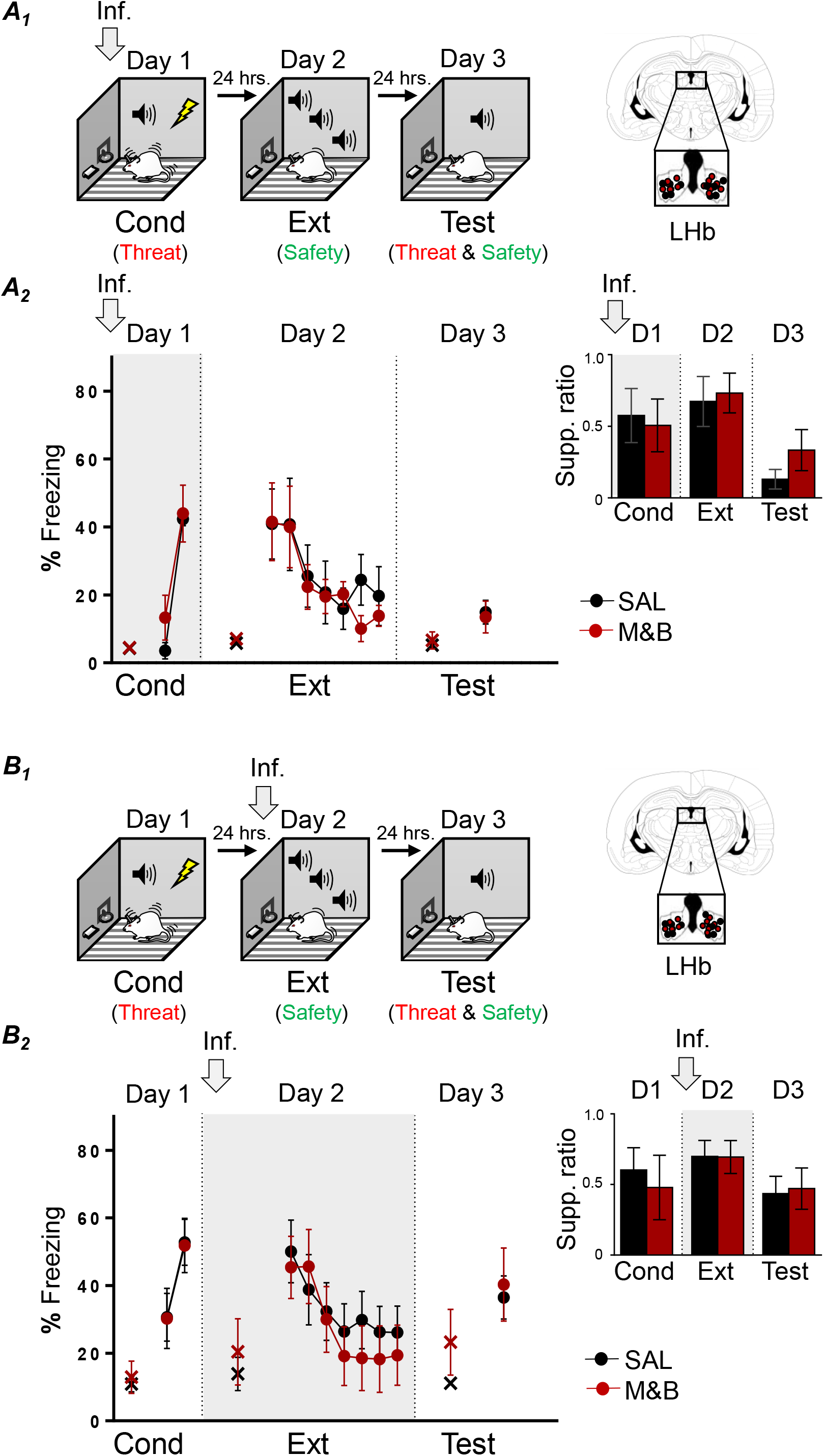
LHb inactivation before threat and extinction training does not affect learning and memory. ***A***_***1***_, ***B***_***1***_, Left, Schematic representation of rats training. Rats were infused with saline solution (SAL) or muscimol and baclofen (M&B) into LHb (gray arrow) before being trained to associate a tone with foot-shocks (day 1; Cond, conditioning). This training represents threat learning (red). One day later, rats were trained to associate the same tone with the absence of foot-shocks (day 2; Ext, extinction), which represents safety learning (green). Finally, the next day, rats were presented with tones only (day 3, Test), which involves the retrieval competition of coexisting threat (red) and safety (green) memories to control defensive responses. Right, Coronal drawings of rat brain atlas showing placements of the injector tips in LHb. ***A***_***2***_ LHb inactivation before threat conditioning (day 1; gray arrow and grayed area in graph), did not affect threat learning (day 1) or its memory formation (day 2), as indicated similar levels of tone-elicited freezing and lever-press suppression ratio (inset: late conditioning (last two-tone trial block), early extinction (first two-tone trial block) and test) between groups (SAL, n = 5; M&B, n = 7). ***B***_***2***_ Infusion of M&B into LHb before extinction training (day 2; gray arrow and grayed area in graph), did not affect, expression of threat memory (day 2), safety learning (day 2) or its memory formation (day 3), as indicated by similar levels of percent freezing and lever-press suppression ratio (inset: late conditioning (last two-tone trial block), early extinction (first two-tone trial block) and test) between groups (SAL, n = 11; M&B, n = 9). Data are shown as ± SEM in blocks of two trials. LHb, lateral habenula. Inf, infusion. Supp. ratio, suppression ratio; x, baseline (pretone) freezing levels.

We next investigated the necessity of LHb on threat memory retrieval as well as on extinction learning and memory by pharmacologically inactivating the LHb before auditory threat extinction training (**Fig. 1**, ***B***_***1***_). Rats were infused with either SAL or M&B into the LHb on day 2, before threat extinction training. One day later (day 3), rats were tested for retrieval of emotional memory to the tone. We found that inactivation of LHb before threat extinction did not affect retrieval of threat memory or extinction acquisition (**Fig. 1**, ***B***_***2***_), as indicated by similar levels of defensive responses across groups in the first block of two-tone trials of extinction training (day 2, freezing: *t*_(18)_ = −0.35, *p* = 0.72; lever-press suppression ratio: *t*_(18)_ = −0.02, *p* = 0.97) and overall during extinction training (day 2, freezing: *F*_(6,108)_ = 0.92, *p* = 0.47; lever-press suppression ratio: *F*_(6,108)_ = 1.90, *p* = 0.08), respectively. Additionally, consistent with a previous report (Song et al., 2017), we found that LHb inactivation before extinction training, did not affect retrieval of extinction-mediated safety memory one day later, as indicated by similar levels of defensive responses across groups during memory test (day 3, freezing: *t*_(18)_ = 0.31, *p* = 0.75; lever-press suppression ratio: *t*_(18)_ = 0.18, *p* = 0.85). Thus, these results suggest that LHb is not necessary for threat memory retrieval as well as not necessary for extinction learning and memory. Together our findings indicate that LHb is not necessary for threat and extinction learning and memory.

### LHb inactivation impairs defensive responses during extinction-dependent retrieval

To evaluate the necessity of LHb in retrieval events after extinction learning, involving the competition of coexisting conditioned-mediated threat memory and extinction-mediated safety memory, we pharmacologically inactivated the LHb before retrieval memory test on day 3 (**Fig. 2**, ***A***_***1***_). Surprisingly, we found that LHb inactivation before memory test, after extinction, abolished defensive responses (freezing and pressing) to preconditioning levels of the experimental group as compared to control group during memory test (day 3, freezing: SAL: 35.59%: M&B: 8.39%; *t*_(14)_ = −3.06, *p* = 0.008; lever-press suppression ratio: SAL: 0.41; M&B: −0.04; *t*_(14)_ = −3.22, *p* = 0.006). Based on a previous LHb inactivation study (Stopper and Floresco, 2014) and to further examine the neuroanatomical specificity of the inactivation effect, after histology, data was separated into three groups: one SAL-infused group and two M&B-infused groups. One group represents, a neuroanatomical control, in which placements missed (Miss M&B) the targeted brain region (placements included dorsal hippocampus and third ventricle) and another group represents rats where placements directly hit (Hit M&B) only in the LHb (**Fig. 2**, ***A***_***2***_; day 3, freezing: SAL: 35.59%, Miss M&B: 37.64%, Hit M&B: 8.39%; *F*_(2,21)_ = 3.54, *p* = 0.04; post hoc comparisons: SAL and Hit M&B: *p* = 0.04, Miss M&B and Hit M&B: *p* = 0.02, and SAL and Miss M&B: *p* = 0.87; lever-press suppression ratio: SAL: 0.41, Miss M&B: 0.39, Hit M&B: −0.04; *F*(2,21) = 5.28, *p* = 0.01; post hoc comparisons: SAL and Hit M&B: *p* = 0.01, Miss M&B and Hit Miss: *p* = 0.01, and SAL and Miss M&B: *p* = 0.94). Results after comparison between experimental (Hit M&B) and control groups (SAL and Miss M&B) reveal that the effects of inactivation on memory retrieval after extinction were circumscribed to the LHb, but not nearby regions. That LHb inactivation before the retrieval test decreased defensive responses could be interpreted as an impairment in the expression of defensive responses. However, this is unlikely given that LHb inactivation did not impair defensive response expression when performed before conditioning or before extinction training (see day 1 in **Fig. 1**, ***A***_***2***_ and day 2 in **Fig. 1**, ***B***_***2***_). Another possible interpretation to this result is that LHb inactivation before testing impaired threat memory retrieval. However, this is also unlikely given that LHb inactivation did not impair threat memory retrieval when performed before extinction training (see first two-tone trial block in day 2 in **Fig. 1**, ***B***_***2***_). Yet another possibility is that LHb inactivation before the retrieval test facilitated extinction memory thereby biasing the expression of the extinction memory over threat memory. This last interpretation suggests that LHb may be necessary to regulate competing threat and safety memories by promoting defensive responses. But before testing this idea further, we decided to test whether the observed LHb inactivation effect was dependent or not on extinction triggered by the tone (rather than the mere passing of time or context exposure) and also evaluate whether LHb inactivation causes unspecific effects on basic locomotor activity or independent motivated behaviors.

**Figure 2.**
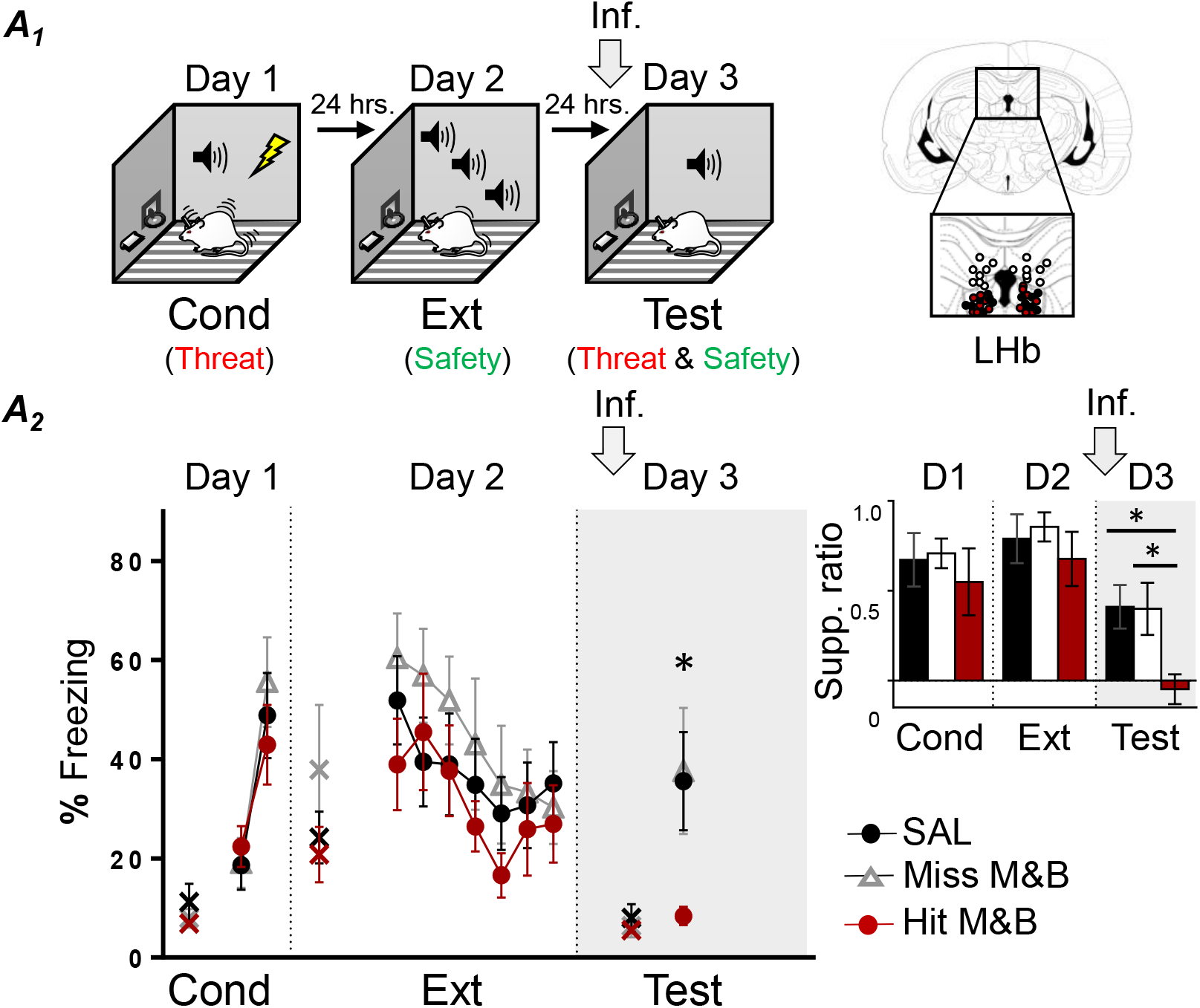
LHb inactivation before memory test, after extinction, abolishes defensive responses. ***A***_***1***_ Left, Schematic representation of rats training. Rats were trained to associate tones with foot-shocks (day 1; Cond, conditioning), which represents threat learning (red). One day later, rats were trained to associate the same tone with the absence of foot-shocks (day 2; Ext, extinction), which represents safety learning (green). Finally, the next day, rats were infused with saline solution (SAL) or muscimol and baclofen (M&B) into LHb (gray arrow) before being presented with tones only (day 3, Test), which involves the retrieval competition of coexisting threat (red) and safety (green) memories to control defensive responses. Right, Coronal drawings of rat brain atlas showing placements of the injector tips in LHb. ***A***_***2***_ LHb inactivation before retrieval test (day 3), after extinction (gray arrow and grayed area in graph), decreased defensive responses, as indicated by lower levels of tone-elicited percent freezing and lever-press suppression ratio (inset: late conditioning (last two-tone trial block), early extinction (first two-tone trial block) and test) in the experimental group as compared to the control groups (SAL, n = 7; Miss M&B, n = 8; Hit M&B n = 9). Data are shown as ± SEM in blocks of two trials. LHb, lateral habenula. Inf, infusion. Supp. ratio, suppression ratio; x, baseline (pretone) freezing levels. *p<0.05.

To evaluate the necessity of LHb to memory retrieval without tone-associated extinction, we inactivated LHb before memory test in rats that were not exposed to the tone during extinction (No-Ext) (**Fig. 3**, ***A***_***1***_). Similar to the experiment above, on day 1, rats were threat conditioned to a tone. The next day (day 2), rats were put back in the conditioned chamber without tone exposure. Finally, on day 3, rats were infused with either SAL or M&B into the LHb and tested for memory retrieval to the tone. We found that LHb inactivation before memory test without tone extinction, did not affect retrieval of conditioned tone threat memory (**Fig. 3 *A***_***1***_), as indicated by similar levels of defensive responses across groups during retrieval test (day 3, freezing: *t*(12) = −0.76, *p* = 0.45; lever-press suppression ratio: *t*_(12)_ = −1.51, *p* = 0.15). This result is consistent with the interpretation that LHb inactivation effect during retrieval test is not due to impairment on the expression of defensive responses or threat memory retrieval to the conditioned tone and that such effect is extinction-dependent to the tone and not the mere passage of time.

**Figure 3.**
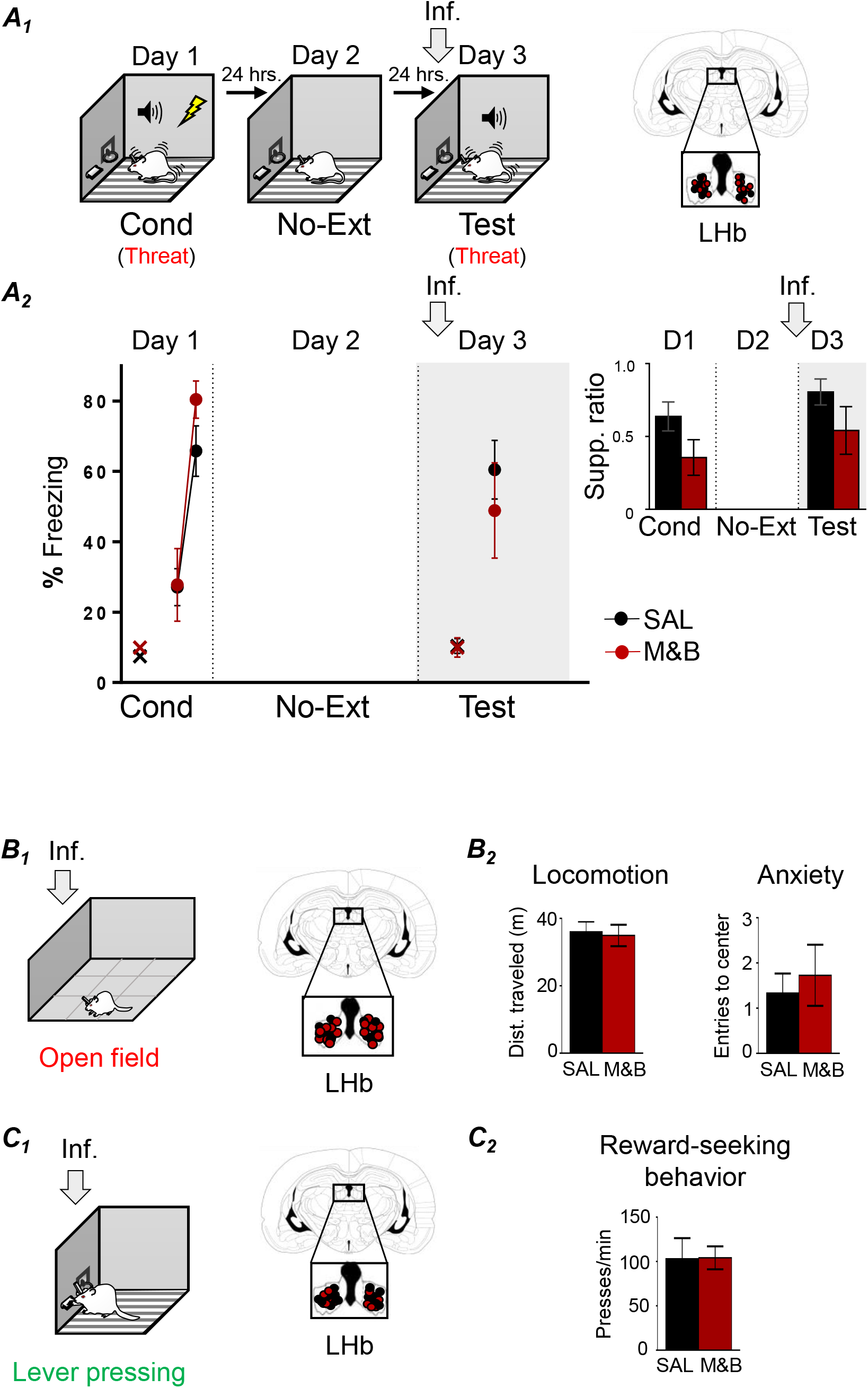
LHb inactivation effect is extinction-dependent and not attributable to impairments in locomotion or other emotional behaviors. ***A***_***1***_ Left, Schematic representation of rats training. Rats were trained to associate a tone with foot-shocks (day 1; Cond, conditioning), which represents threat learning (red). One day later, rats were put back in the conditioning chamber (day 2; No Ext, no tone extinction). The next day, rats were infused with saline solution (SAL) or muscimol and baclofen (M&B) into LHb (gray arrow) before being presented with tones only (day 3, Test), which represents the retrieval of the threat-related memory (red). Right, Coronal drawings of rat brain atlas showing placements of the injector tips in LHb. ***A***_***2***_ LHb inactivation before retrieval test (day 3) without extinction (gray arrow and grayed area in graph), did not affect defensive responses to the tone, as indicated by similar levels of tone-elicited percent freezing and lever-press suppression ratio (inset: late conditioning (last two-tone trial block and test) between control and experimental groups (SAL, n = 8; M&B, n = 6). Data are shown as ± SEM in blocks of two trials. Supp. ratio, suppression ratio; x, baseline (pretone) freezing levels. ***B***_***1***_ Left, Schematic representation of open field task. Right, Coronal drawings of rat brain atlas showing the placements of the injector tips in LHb. ***B***_***2***_ LHb inactivation did not affect locomotion or anxiety-related behavior, as indicated by similar distance traveled and number of entries to the center of the field (SAL, n = 12; M&B n = 11). ***C***_***1***_ Left, Schematic representation of lever pressing. Right, Coronal drawings of rat brain atlas showing the placements of the injector tips in LHb. ***C***_***2***_ LHb inactivation did not affect reward-seeking behavior, as indicated by similar presses per minute to obtain food (SAL, n = 5; M&B n = 7). LHb, lateral habenula. Inf, infusion.

Next, in a different group of animals, we tested the effect of LHb inactivation on basic locomotion activity, an innate aversively motivated behavior (exploration in an open field) and an appetitively motivated behavior. We tested this by infusing SAL or M&B into the LHb before an open field task and before a lever pressing test. We found that LHb does not affect locomotion, anxiety-related (**Fig. 3**, ***B***_***1***_ **and *B***_***2***_) or reward-seeking behaviors (**Fig. 3**, ***C***_***1***_ ***C***_***2***_), as indicated by similar levels of total distance traveled (*t*_(21)_ = −0.25, *p* = 0.80) and entries to the center of the open field (*t*_(21)_ = 0.49, *p* = 0.622), as well as similar levels of presses per minute to obtain food (*t*_(10)_ = 0.04, *p* = 0.96), respectively. These control experiments suggest that LHb inactivation (on its own) does not affect general movement, hunger, motivation to explore related to anxiety like-behavior or reward-seeking behavior.

### LHb inactivation impairs defensive responses during latent inhibition retrieval

Together, our above results are consistent with the idea that rather than being necessary for simple task performance (freezing or not to the tone), LHb-mediated activity is crucial for flexible and adaptive behaviors during more complex situations in mammals (Baker et al., 2017). Threat and extinction learning and memory as well as expression of defensive responses by themselves may represent relatively simple situations as compared to flexibly selecting the adaptive emotional memory (threat or safety) to control (promote or suppress) the expression of defensive behaviors. Such situation occurs when coexisting threat and safety memories compete for control of behavior after extinction training. To further test the idea that LHb is necessary in conditions where coexisting threat and safety memories compete for control of behavior, we switched the order of opposing emotional learning experiences (safety learning followed by threat learning rather than threat learning followed by safety learning) by using latent inhibition (**Fig. 4**, ***A***_***1***_). To do this, on day 1, rats were either trained to acquire a latent inhibition-mediated safety memory by presenting tones without foot-shocks (latent inhibition group: LI) or simply placed in the conditioning chamber (no tones or foot-shocks; No latent inhibition group: No-LI). On day 2, both group of rats were threat conditioned to the tone. One day later (day 3), rats were tested for memory retrieval to the tone. As expected, we found that pre-exposure to a tone, weakened subsequent threat conditioning to tones (**Fig. 4**, ***A***_***2***_), as indicated by lower defensive response levels of the LI group with respect to No-LI group at the beginning of conditioning (first two-tone trial block; freezing: No-LI: 22.30%; LI: 9.03%; *t*(21) = −2.16, *p* = 0.04; lever-press suppression ratio: No-LI: 0.32; LI: −0.15; *t*(21) = −2.64, *p* = 0.01). On day 3, both groups showed similar defensive response levels (freezing: *t*(21) = −0.33, *p* = 0.74; lever-press suppression ratio: *t*(21) = −0.98, *p* = 0.33). This result suggests that cued-mediated latent inhibition leads to the formation of a safety memory that retards conditioning learning to the tone. Next, to evaluate the necessity of LHb during the competition of coexisting latent inhibition-mediated safety memory and conditioned-mediated threat memory, we pharmacologically inactivated the LHb before a post-conditioning memory test (**Fig. 4**, ***B***_***1***_). On day 1, rats were pre-exposed to the tone. The next day (day 2), rats were conditioned to the tone by paring tones with foot-shocks.

**Figure 4.**
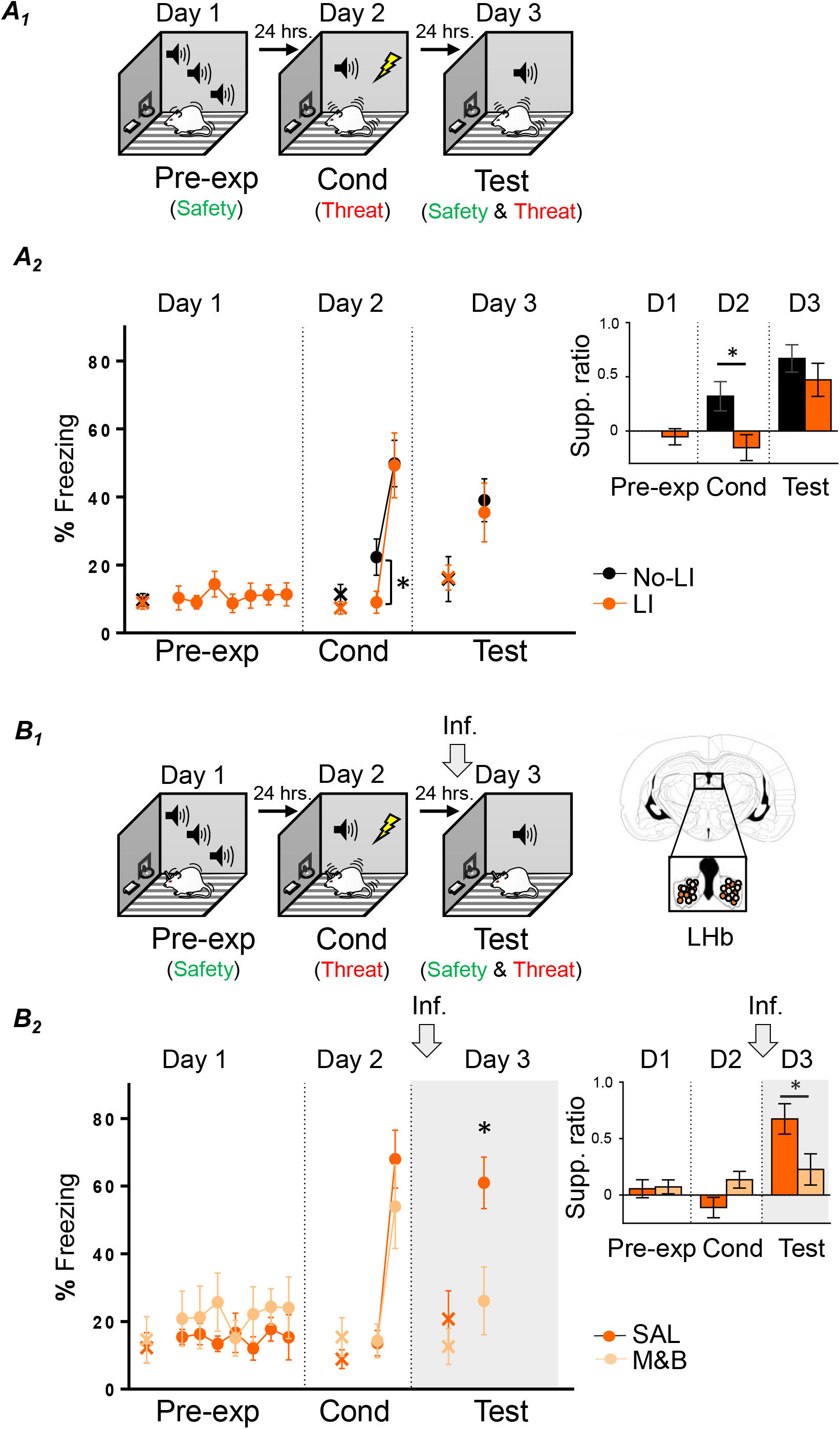
LHb inactivation before retrieval test, after conditioned latent inhibition, abolishes defensive responses. ***A***_***1***_, Schematic representation of rats training. Rats were either presented with tones (latent inhibition group: LI), which represents safety learning (green), or simply placed in the conditioning chamber (No-latent inhibition group: No-LI; day 1; Pre-exp, pre-exposure). One day later, both groups were placed in the same behavioral chamber and trained to associate a tone with foot-shocks (day 2; Cond, conditioning), which represents threat learning (red). Finally, the next day, were presented with tones only (day 3, Test), which involves the retrieval competition of coexisting safety (green) and threat (red) memories to control defensive responses. ***A***_***2***_, Tone pre-exposure weakened subsequent threat learning, as indicated by lower tone-elicited percent freezing and lever-press suppression ratio (inset: late pre-exposure, early conditioning and test) levels of the LI group, as compared to No-LI group at the beginning of conditioning (No-LI, n = 11; LI, n = 12). ***B***_***1***_, Left, Schematic representation of rats training. Rats were pre exposed to tones alone (day 1; Pre-exp, pre exposure), which represents safety learning (green). One day later, rats were trained to associate the tone with foot-shocks (day 2; Cond, conditioning), which represents threat learning (red). Finally, the next day, rats were infused with saline solution (SAL) or muscimol and baclofen (M&B) into LHb (gray arrow) before being presented with tones only (day 3, Test), which involves the retrieval competition of coexisting safety (green) and threat (red) memories to control defensive responses. Right, Coronal drawings of rat brain atlas showing placements of the injector tips in LHb. ***B***_***2***_, LHb inactivation before retrieval test (day 3), after latent inhibition and conditioning (gray arrow and grayed area in graph), decreased defensive responses as indicated by lower levels of tone-elicited percent freezing and lever-press suppression ratio (inset: late conditioning (last two-tone trial block), early extinction (first block of trails) and test) in the experimental group as compared to the control group (SAL, n = 8; M&B, n = 5). Data are shown as ± SEM in blocks of two trials. LHb, lateral habenula. Inf, infusion. Supp. ratio, suppression ratio; x, baseline (pretone) freezing levels. *p<0.05.

Finally, on day 3, rats were infused with either SAL or M&B into the LHb and tested for memory retrieval to the tone. Similar to infusion during retrieval after extinction, we found that LHb inactivation before memory test, after conditioning, decreased defensive response levels of the experimental group as compared to control group during memory test, (**Fig. 4**, ***B***_***2***_; day 3, freezing: SAL: 61.00%, M&B: 26.09%; *t*_(11)_ = −2.80, *p* = 0.01; lever-press suppression ratio: SAL: 0.65%, M&B: 0.22%; *t*_(11)_ = −2.25, *p* = 0.04). Since we ruled out above that LHb inactivation impairs the expression of defensive responses and threat memory retrieval, this finding supports the idea that LHb inactivation before the retrieval test biased toward facilitating the expression of the latent inhibition-mediated safety memory over the conditioned-mediated threat memory. This last interpretation suggests that, independently of when those opposing emotional memories are formed (safety before threat memory or threat before safety memory), LHb is necessary to regulate competing and coexisting threat and safety memories by promoting defensive responses to ultimately exert control over adaptive behaviors.

### LHb inactivation switches choice behavior bias from avoidance to reward-seeking

Besides influencing adaptive defensive responses triggered by threats like freezing (this study and (Martinez-Canabal et al., 2019)) and avoidance (Thornton and Bradbury, 1989; Tomaiuolo et al., 2014; Trusel et al., 2019), LHb activity has been recently implicated in biasing choice behavior (Stopper and Floresco, 2014; Baker et al., 2017; Mathis et al., 2017). Therefore, using platform-mediated avoidance (PMA) task (Bravo-Rivera et al., 2014; Diehl et al., 2019; Bravo-Rivera and Sotres-Bayon, 2020), we tested the effect of LHb inactivation when rats are challenged to choose between taking action to avoid a threat or taking action to approach food guided by competing coexisting threat and safety memories. After trained to press a lever to obtain food, rats were trained for 10 days to learn to avoid a threat (foot-shock) signaled by a tone, by stepping into the safe platform (**Fig. 5A**). During early PMA training (day 2), rats acquire classical conditioning (tone-shock threat association), but barely any instrumental learning (tone-platform safety association), as indicated by moderate levels of freezing and relatively low levels of avoidance responses (**Fig. 5B**; *t*_(28)_ = 0.42, *p* = 0.67), respectively. In contrast, by late PMA (day 10), freezing responses remained similar to early PMA (34.49% on day 2 vs. 44.71% on day 10; *t*_*(28)*_ = −1.56, *p* = 0.12), but avoidance responses dramatically increased (30.00% on day 2 vs. 83.33% on day 10; *t*_(28)_ = −4.67, *p* < 0.00006), as indicated by moderate levels of freezing and high levels of avoidance responses (freezing: 44.71% and avoidance: 83.333%; *t*_(28)_ = −4.95, *p* < 0.000006). These results suggest the formation of a tone threat memory during early PMA, whereas the establishment of a tone safety memory during late PMA. Finally, the next day (day 11), rats were infused with either SAL or M&B into the LHb and tested for choice behavior (avoid or approach to obtain food) guided by competing threat and safety memories. Notably, we found that LHb inactivation before PMA retrieval test, impaired defensive responses, as indicated by decreased freezing and avoidance responses of the experimental group as compared to control group during PMA test (**Fig. 5C**: freezing: SAL: 29.85%, M&B: 5.22%; *t*_(13)_ = −2.73, *p* = 0.01; **Fig. 5D**_***1***_: avoidance SAL: 59.26%, M&B: 15.39%; *t*_(13)_— = −3.00, *p* = 0.01). Although showing a similar decrease trend, averaged lever press suppression ratio during test was not significantly different after inactivation (**Fig. 5C inset**: SAL: 0.63%, M&B: 0.25%; (*t*_(13)_ = 1.85, *p* = 0.08). Yet, analyzing the percent time spent in platform during the tone and suppression ratio (in 6 sec bins) revealed that LHb inactivation blocked both avoidance and suppression ratio (i.e. inactivation facilitated lever press suppressing during tone compared to pretone) compared to animals in the control group (**Fig. 5D**_***2***_: time in platform *F*_(1,13)_=7.50, *p* = 0.01; post hoc comparisons: −6 to −1 s, *p* = 0.71; 0-6 s, *p* = 0.23; 7-12 s, *p* = 0.0048; 13-18 s, *p* = 0.0008; 19-24 s, *p* = 0.007; 25-30 s, *p* = 0.02. Inset, suppression ratio *F*_(1,13)_ = 6.55, *p* = 0.02; post hoc comparisons: 0-6 s, *p* = 0.32; 7-12 s, *p* = 0.11; 13-18 s, *p* = 0.0042, 19-24 s, *p* = 0.03; 25-30 s, *p* = 0.44). In sum, LHb inactivation biased the ability to actively choose between avoiding a threat or approaching food, towards driving reward-seeking behavior, suggesting that LHb is necessary to bias choice towards the promotion of defensive responses when competing threat and safety memories guide motivated behavior.

**Figure 5.**
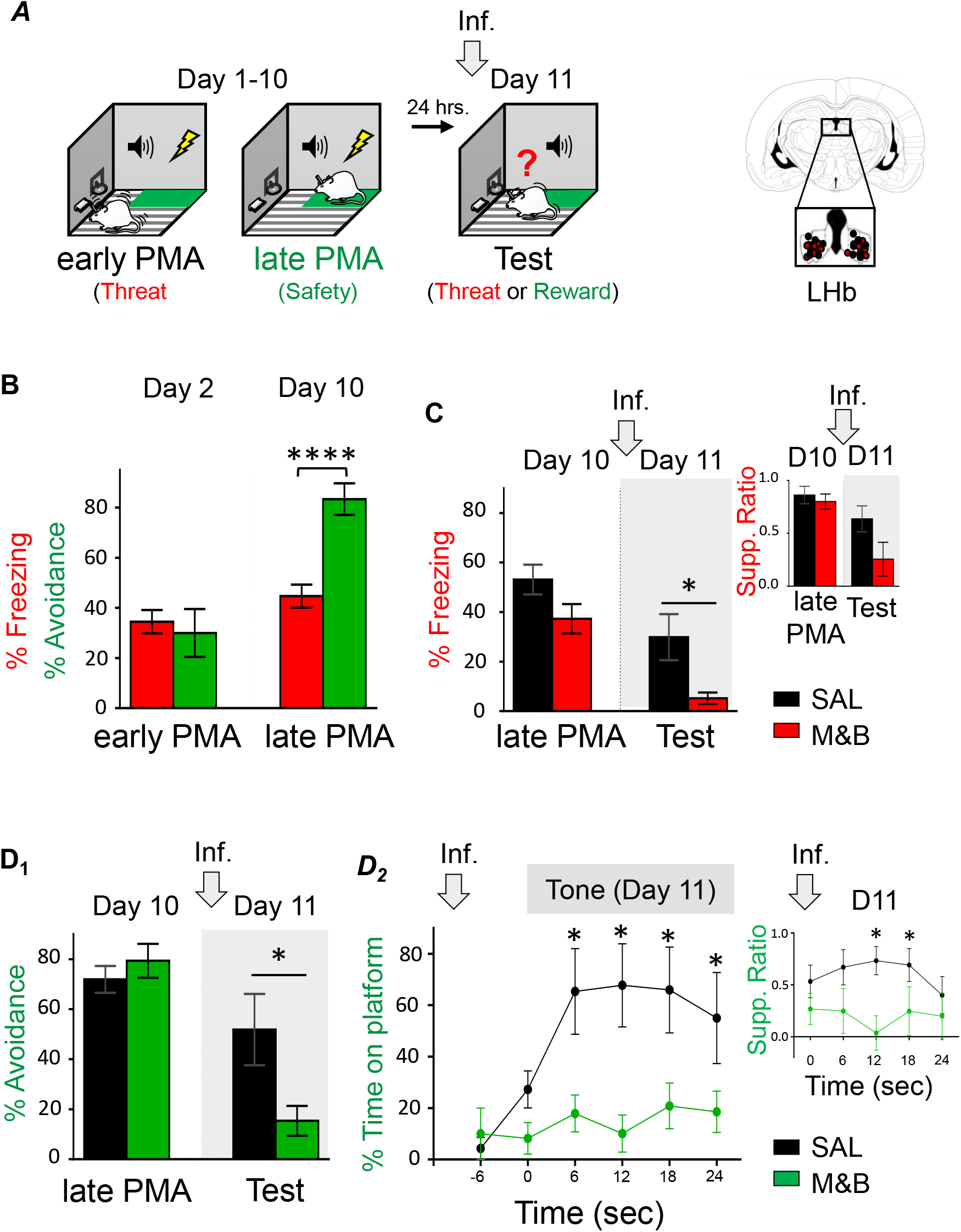
LHb inactivation before platform-mediated avoidance retrieval test biases choice-mediated defensive behavior towards reward-seeking. ***A***, Left, Schematic representation of rats training. Rats were trained to associate tone with foot-shocks as well as associating the tone with moving to a safety platform learn to avoid the shock (day 1-10; platform-mediated avoidance (PMA)). Early PMA (day 2) represents classical conditioned-mediated threat learning (red), whereas late PMA (day 10) represents instrumental avoidance-mediated safety learning (green). One day later, rats were infused with saline solution (SAL) or muscimol and baclofen (M&B) into LHb (gray arrow) before being presented with tones only (day 11, Test), which involves the retrieval competition of coexisting threat (red) and safety (green) memories to control choice behavior. Right, Coronal drawings of rat brain atlas showing placements of the injector tips in LHb. ***B,*** During early PMA training (day 2), freezing and avoidance levels are similar. In contrast, during late PMA training (day 10), avoidance levels are higher than freezing levels. Thus, avoidance levels dramatically increased with PMA training while freezing levels remain similar (n = 15). ***C***, LHb inactivation before PMA test (gray arrow and grayed area in graph), impaired retrieval of PMA memory, as indicated by lower tone-elicited percent freezing responses (but not suppression ratio) of the experimental group as compared to control group (SAL n = 7; M&B = 8). ***D,*** LHb inactivation before PMA test (gray arrow) biased choice-mediated avoidance towards reward-seeking behavior as indicated by lower tone-elicited percent avoidance (***D***_***1***_) and time spent in platform (***D***_***2***_), as well as lower suppression ratio (inset) in the experimental group as compared to the control group (SAL n = 7; M&B = 8). Data are shown as ± SEM in blocks of two trials in B, C and D_1_ and in 6 s bins in D_2_. LHb, lateral habenula. Inf, infusion. Supp. ratio, suppression ratio; **** p<0.00001; *p<0.05.

## Discussion

We investigated the conditions under which LHb is necessary to regulate defensive responses to a threat. By combining pharmacological inactivations with various behavioral paradigms in rats, including tasks that involve retrieval of competing threat and safety memories, our study reveals a previously unknown role for the LHb in driving reactive and active defensive responses during emotional conflict. These findings add to a growing body of evidence showing that LHb is involved in the regulation of aversively motivated behaviors and highlight emotional conflict as a condition under which this brain region is critical to promote defensive responses.

### LHb promotes defensive strategies during threat/safety memory conflict

Even after successful extinction learning to threat, conditioned defensive responses often return with the simple passage of time (Rescorla, 2004; Sotres-Bayon et al., 2006); even after a single day (Quirk, 2002). By using c-Fos as neural activity marker, a previous study identified that high LHb activity is positively correlated with high threat recovery after extinction (Martinez-Canabal et al., 2019). Consistently, in this study, we found that LHb inactivation prevents the recovery of extinguished defensive responses. Together, these findings indicate that LHb is not only recruited but is necessary to prevent the expression of the extinction memory thereby allowing defensive responses to return after extinction.

LHb activity may normally prevent the retrieval of safety memories in the service of driving defensive responses by suppressing downstream structures involved in appetitive motivated behaviors. A recent study found that optogenetic stimulation of dopaminergic activity in the ventral tegmental area (VTA), a prominent LHb projection site, facilitates extinction memory (Salinas-Hernandez et al., 2018). Another recent study showed that chemogenetic inhibition of serotonergic activity of dorsal raphe receiving LHb input provides animals with resilience during aversive conditions (Varga et al., 2003; Andalman et al., 2019). Because LHb is known to potently inhibit brainstem dopaminergic and serotoninergic activity (Varga et al., 2003; Matsumoto and Hikosaka, 2007; Lammel et al., 2012; Andalman et al., 2019), it is possible that suppressing LHb activity disinhibits these monoaminergic pathways thereby facilitating retrieval of safety memories (extinction and latent inhibition) and resilience in aversive conditions. In line with these findings, we suggest that LHb activity normally prevents expression of the extinction memory possibly by inhibiting VTA dopaminergic activity in the service of signaling the prevailing threat memory during recovery. This interpretation is further consistent with recent findings that indicate that LHb activity increases under aversive conditions, leading individuals towards negative expectations (Shabel et al., 2019). Furthermore, blockade of LHb activity with ketamine results in antidepressant effects (Yang et al., 2018; Cui et al., 2019) and decreased LHb activity mediated by enhanced neurogenesis prevents threat recovery (Martinez-Canabal et al., 2019). Thus, we suggest that when LHb is suppressed (with pharmacological inactivation, ketamine or enhanced neurogenesis), the negative expectation signal (worse than expected or “pessimism”) is blocked (likely by disinhibiting monoaminergic pathways involved in motivation), thereby allowing individuals to be able to explore other options such as approaching food when available.

Extinction allows us to evaluate how coexisting threat and extinction memories compete for the control of defensive responses (Rescorla, 2001; Myers and Davis, 2002; Quirk, 2002; Quirk and Mueller, 2008; Herry et al., 2010; Martinez-Canabal et al., 2019), and our findings suggest that the LHb is a critical brain region necessary to promote defensive responses during retrieval of conflicting threat and extinction memory. This interpretation is consistent with the idea that LHb activity is shaped by previous aversive and appetitive emotional associations (Wang et al., 2017) and that experience-dependent modifications in this brain region are necessary for the selection of defensive responses (Amo et al., 2014). Together, these findings suggest the possibility that LHb is involved in promoting defensive responses in conditions where memory retrieval is guided by coexisting previous aversive and appetite experiences. Another condition that allows us to evaluate the competition of threat and safety memories, although in the inverse order, is following latent inhibition. We found that LHb is necessary to promote defensive responses under both retrieval test task conditions (after extinction and after latent inhibition). These findings suggest that, independently of the order of acquisition of threat and safety learning, LHb is necessary to guide defensive responses mediated by the competition of coexisting emotional experiences (threat vs safety or safety vs threat). Thus, together these results indicate that, LHb may not mediate recovery of extinguished defensive responses per se but rather the promotion of threat-related behaviors in retrieval events involving competition of threat and safety memories.

### LHB promotes defensive responses during conflict-mediated choice behavior

Our finding that LHb is necessary for retrieval of PMA is consistent with previous studies that show that LHb is necessary for avoidance learning (Wilcox et al., 1986; Thornton and Bradbury, 1989). However, PMA differs from traditional avoidance tasks in that it uses choice behavior to evaluate competing emotional memories during conflict (Bravo-Rivera et al., 2014; Diehl et al., 2019; Bravo-Rivera and Sotres-Bayon, 2020). During PMA training, a platform acquires a positive motivational value (safety) while the grid acquires a negative value (threat). Then rats are challenged, during PMA retrieval (no shock delivered), to decide, guided by competing emotional memories signaled by a tone, whether to step onto the safe platform or press a lever to obtain food. Normally, rats choose to avoid the shock by stepping into the platform, however LHb inactivated rats pressed the lever to obtain food despite the threat. This finding suggests that LHb is necessary to promote defensive responses over reward-seeking behavior during conflict choice guided by memories with opposing motivational valences (threat and safety memories).

A previous study reported increase activity of dopaminergic neurons that allow successful avoidance responses (Oleson et al., 2012). The LHb inhibits dopaminergic VTA activity, through GABAergic neurons in the tail of this brain structure (the rostromedial tegmental nucleus), promoting defensive responses, thereby putting a “break” on reward-seeking (Jhou et al., 2009; Quirk and Sotres-Bayon, 2009; Proulx et al., 2018). Our pharmacological inhibition of the LHb may allow disinhibition of VTA dopaminergic activity leading to a facilitation of reward-seeking thereby making the individual more prone to risky behaviors. This is consistent with a previous study showing that LHb inactivation increased hesitation to make a choice between risky reward options associated with different subjective values (Stopper and Floresco, 2014) and the notion that LHb is involved in guiding survival decisions based on stimuli with learned motivation value (Hikosaka, 2010; Baker et al., 2016). Thus, our PMA findings support the notion that LHb is critical for biasing choice behavior mediated by value-based decision making and highlight its fundamental role in promoting defensive behavioral strategies during conflict-mediated choice behavior.

### LHb as a critical node in emotional conflict

Overall, our study reveals that emotional memory conflict is a fundamental survival condition under which LHb is required to promote defensive strategies. This conclusion is consistent with the emerging notion that LHb acts as a critical node in the control of emotional behaviors guided by aversive events in health and disease, at least in part, mediated by regulating downstream midbrain dopaminergic and serotoninergic structures (Hu et al., 2020). We propose that during emotional conflict, LHb activity is required to bias experience-dependent emotional behavior towards defensive behavioral responses and away from reward-seeking behavior, likely by suppressing downstream structures involved in appetitive motivated behaviors. Accordingly, a lack of activity in LHb during emotional conflict results in a shift towards reward-seeking behavior and away from defensive responding, likely through disinhibition of downstream structures involved in appetitive motivated behaviors. Identifying the brain regions embedded in a complex network that are required for conflict-mediated behavioral responses, may be key for understanding how the brain balances information about competing emotional memories to adaptatively regulate emotional behaviors in health and disease.

## Author Contributions

GV-H and FS-B designed research; GV-H performed experiments; GV-H and FS-B analyzed data and wrote the paper.

## Acknowledgements

We thank Ana Peñas-Rincón for technical assistance in avoidance experiments and Gregory J. Quirk, Christian Bravo-Rivera and Leticia Ramirez-Lugo for helpful discussions and comments on the manuscript. This work was supported by Consejo Nacional de Ciencia y Tecnología (CONACyT, grant PN2463), Dirección General de Asuntos del Personal Académico de la Universidad Nacional Autónoma de México (UNAM, grants IN205417 and IN214520) and the International Brain Research Organization (Return Home fellowship) to FS-B. G V-H is a doctoral student at Programa de Doctorado en Ciencias Biomédicas at UNAM, and was supported by a CONACyT fellowship (658352).

